# Prepubertal ovariectomy alters dorsomedial striatum indirect pathway neuron excitability and explore/exploit balance in female mice

**DOI:** 10.1101/2021.06.01.446609

**Authors:** Christopher D. Hall, Linda Wilbrecht, Kristen Delevich

**Affiliations:** Department of Psychology, University of California, Berkeley, CA 94720, USA; Helen Wills Neuroscience Institute, University of California, Berkeley, CA 94720, USA

## Abstract

Decision-making circuits are modulated across life stages (e.g. juvenile, adolescent, or adult)—as well as on the shorter timescale of reproductive cycles in females—to meet changing environmental and physiological demands. Ovarian hormone signaling may contribute to this flexibility by regulating neural circuits involved in adaptive decision making. Here, we examined how prepubertal ovariectomy (pOVX) influences adult performance in an odor-guided multiple-choice reversal task. During reversal learning, pOVX females exhibited a distinct pattern of errors compared to sham-operated controls. To characterize decision strategies, we fit a reinforcement learning model to trial-by-trial behavior across both the initial Discrimination and Reversal phases. Sham females showed a significant increase in the inverse temperature parameter (β) during Reversal, consistent with a shift towards a more exploitative choice policy favoring high-value options. By contrast, pOVX females failed to show a phase-dependent increase in β, suggesting they continued to use a more exploratory strategy. To identify a neural correlate of this behavioral phenotype, we performed whole-cell patch clamp recordings within the dorsomedial striatum (DMS), a region implicated in regulating action selection and explore/exploit choice policy. Intrinsic excitability of dopamine receptor type 2-expressing (D2R) indirect pathway spiny projection neurons (iSPNs) was significantly higher in pOVX females compared to both sham-operated and unmanipulated controls. Finally, to test whether increased D2R(+) iSPN excitability could causally contribute to the exploratory reversal phenotype, we chemogenetically activated DMS D2R(+) neurons in intact female mice. This manipulation increased exploratory choice during reversal learning, recapitulating the pattern observed following pOVX. Together, these data suggest that pubertal status may influence how females adapt their decision strategy between Discrimination and Reversal phases, potentially through long-lasting modulation of DMS iSPN intrinsic excitability.

## Introduction

As animals interact with their environment in pursuit of rewards in the form of food, water, mates etc., they learn from trial and error to guide their future choices. This process involves learning from positive and negative feedback and also, importantly, deciding how learned information should influence choice, referred to as choice policy. Reinforcement learning (RL) models (Sutton and Barto, 2020) have provided a useful framework for understanding and quantifying aspects of trial-and-error learning, including choice policy. A classic RL problem that hinges on choice policy is the explore/exploit tradeoff. If an animal (or any agent for that matter) adopts an exploit policy, it will consistently select the highest estimated value option but may miss out on better alternative options. On the other hand, if an animal favors a more exploratory choice policy, characterized by less value-dependent, more stochastic choice behavior, it may discover new and better options more readily (Daw et al., 2006; Sutton and Barto, 2020). Importantly, the optimal balance of exploration and exploitation may depend on the statistics of the environment and/or the needs of the animal as defined by its particular physiological or developmental state (Addicott et al., 2017; Cohen et al., 2007; Frank et al., 2009; Gopnik, 2020; Humphreys et al., 2015; Lenow et al., 2017; Wilbrecht and Davidow, 2024). In humans, choice behavior generally becomes less exploratory and more exploitative during the transition from childhood to adulthood (Eckstein et al., 2022; Gopnik, 2020; Nussenbaum and Hartley, 2019; Xia et al., 2021). Natural fluctuations in ovarian hormones across the estrous cycle or exogenous estradiol administration have been shown to regulate aspects of value-based decision making in female rodents (Golden et al., 2025; Orsini et al., 2021; Uban et al., 2012), including explore/exploit balance (Verharen et al., 2019b). These data suggest that the rise in ovarian hormones at puberty could contribute to the developmental shift in choice policy during adolescence in females. In previous work, we observed that pOVX altered performance in a multiple choice reversal task in adult C57BL/6 mice (Delevich et al., 2020a). Compared to intact females, pOVX females showed lower ratios of perseverative to regressive errors during reversal learning, but the potential underlying biological processes that contributed to this behavioral effect remained unclear.

The DMS is implicated in the regulation of goal-directed action selection (Balleine et al., 2021; Matamales et al., 2020; Nonomura et al., 2018; Peak et al., 2020; Tai et al., 2012) and choice policy (Collins and Frank, 2014), and previous work suggests that enhancing the activity of D2R(+) SPNs in the dorsal striatum biases choice behavior to be more exploratory (Delevich et al., 2022; Lee et al., 2015) but see (Verharen et al., 2019a). While nuclear estrogen receptors are notably absent from the dorsal striatum in adulthood (Krentzel et al., 2021), extranuclear estrogen receptors (ERα, ERβ, and GPER1) localize to SPNs, glia, and the presynaptic terminals of striatal GABAergic and cholinergic interneurons of adult female rats (Almey et al., 2012). At the neuronal level, estrous cycle has been shown to regulate the intrinsic excitability of SPNs located within the rodent striatum (Alonso-Caraballo and Ferrario, 2019; Proaño et al., 2018). Studies examining the influence of estrous cycle on SPN physiology have been primarily performed in rats, where SPN cell types were not distinguished, but see (Tansey et al., 1983). Taken together, these findings raise the question of whether pubertal status influences choice strategies employed by females by modulating striatal SPN physiology. Here we focused on D2R(+) SPNs of the indirect pathway (iSPNs) within the DMS, whose activity we hypothesized regulates explore/exploit balance in decision making based on theoretical predictions (Collins and Frank, 2014; Dunovan and Verstynen, 2016) plus genetic (Beeler et al., 2010; Kwak et al., 2014) and pharmacological (Lee et al., 2015; McCoy et al., 2019) evidence.

In the current study, after analyzing raw behavioral data, we applied RL modeling to examine how pOVX influenced learning and choice policy in the odor-based reversal learning task. We next examined the influence of pOVX on the intrinsic excitability of genetically identified D2R(+) SPNs within the DMS of adult female mice. Finally, we chemogenetically activated D2R(+) neurons within the DMS of female mice during reversal learning and applied an RL model to determine whether this manipulation recapitulated the choice policy phenotype observed in pOVX females. We found that intact adult females flexibly adjusted their choice policy across task phases, shifting from a relatively exploratory policy during Discrimination to a more exploitative policy during Reversal, as captured by a significant increase in the inverse temperature parameter (β) across phases. In contrast, pOVX females did not demonstrate this task-phase-dependent shift, retaining a comparatively exploratory choice policy during Reversal. These behavioral differences were accompanied by increased intrinsic excitability of D2R(+) SPNs in the DMS of pOVX compared to sham-operated females. Finally, chemogenetic activation of D2R(+) SPNs within the DMS promoted a more exploratory choice strategy during reversal learning in intact female mice, resembling pOVX female behavior. Together, these data suggest that pubertal status influences how female mice flexibly adjust their choice strategy during value-based decision making, via the modulation of D2R(+) SPN activity in the DMS.

## Materials & Methods

### Animals

Female C57BL/6NCR (Charles River), Drd2-eGFP BAC (GENSAT), and D2-Cre ER43 (MMRC) mice were bred in-house. Drd2-eGFP BAC and D2-Cre ER43 mice were bred onto the C57BL/6NCR background for at least 5 generations. All mice were weaned on postnatal day (P)21 and housed in groups of 2-3 same-sex siblings on a 12:12 hr reversed light:dark cycle (lights on at 2200 h). All procedures were approved by the Animal Care and Use Committee of the University of California, Berkeley and conformed to principles outlined by the NIH Guide for the Care and Use of Laboratory Animals.

### Prepubertal Ovariectomy

Prepubertal ovariectomy was performed as previously described (Delevich et al., 2020b; Klappenbach et al., 2023). To eliminate ovarian hormone exposure during and after puberty, ovariectomies were performed before puberty onset at P25. Prior to ovariectomy (OVX) surgery, all female mice were visually inspected to confirm that vaginal opening had not occurred. Prior to surgery, mice were injected with 0.05 mg/kg buprenorphine and 10 mg/kg meloxicam subcutaneously and were anesthetized with 1–2% isoflurane during surgery. The incision area was shaved and scrubbed with ethanol and betadine. Ophthalmic ointment was placed over the eyes to prevent drying. A 1 cm incision was made with a scalpel in the lower abdomen across the midline to access the abdominal cavity. The ovaries were clamped off from the uterine horn, with locking forceps and ligated with sterile sutures. After ligation, the ovaries were excised with a scalpel. The muscle and skin layers were sutured, and wound clips were placed over the incision for 7-10 days to allow the incision to heal. An additional injection of 10 mg/kg meloxicam was given 24 and 48 h after surgery. Sham control surgeries were performed in which fat pads were visualized but the ovaries were not clamped, ligated, or excised. Mice were allowed to recover on a heating pad until ambulatory and were post-surgically monitored for 7–10 days to check for normal weight gain and signs of discomfort/distress. Mice were co-housed with 1–2 siblings who received the same surgical treatment. To confirm the success of prepubertal ovariectomies, necropsy was performed on a subset of adult sham and ovariectomized mice to confirm that the uteri of pOVX mice were underdeveloped compared to age-matched sham females (data not shown).

### 4 choice odor-based reversal task

Sham or pOVX mice were tested in an odor-based reversal task that has previously been described in detail (Johnson C, 2011; Johnson et al., 2016) as young adults (P60–P70). The task is designed such that only the odor cue is predictive of reward, while spatial and egocentric information is irrelevant. Briefly, mice were food restricted to ∼85% body weight by the Discrimination phase. Mice were habituated to the testing arena on day 1, they were taught to dig for a honey nut cheerio reward in a pot filled with unscented wood shavings on day 2, underwent a 4 choice odor Discrimination on day 3, and finally, were tested on Recall of the previously learned odor-reward association, which was immediately followed by a Reversal phase on day 4. During the Discrimination phase of the task, mice learned to discriminate among four pots with different scented wood shavings (anise, clove, litsea, and thyme). All four pots were sham-baited with cheerio (under wire mesh at bottom) but only one pot was rewarded (anise). The pots of scented shavings were placed in each corner of an acrylic arena (12”, 12”, 9”) which was divided into four quadrants. Mice were placed in a cylinder in the center of the arena, and a trial started when the cylinder was lifted. Mice were then free to explore the arena and indicate their choice by making a bi-manual dig in one of the four pots of wood shavings. The cylinder was lowered as soon as a choice was made. If the choice was incorrect, the trial was terminated and the mouse was gently encouraged back into the start cylinder. Trials in which no choice was made within 3 minutes were considered omissions. If mice omitted for two consecutive trials, they received a reminder: a baited pot of unscented wood shavings was placed in the center cylinder and mice dug for the “free” reward. Mice were disqualified if they committed four pairs of omissions. The location of the four odor scented pots was shuffled on each trial, and criterion was met when the mouse completed 8 out of 10 consecutive trials correctly. 24 hours after completing Discrimination, mice were tested for Recall of the initial odor Discrimination to criterion, after which, mice immediately proceeded to the Reversal phase in which the previously rewarded odor (anise) was no longer rewarded, and a previously unrewarded odor (clove) was now rewarded. During the Reversal phase, Odor 4 (thyme) was replaced by a novel odor (eucalyptus) that was unrewarded. Again, mice were run until they reached a criterion of 8 out of 10 consecutive correct trials.

### 4 choice odor-based reversal task analysis

To compare reversal task performance across groups, trials to criterion and errors (incorrect choices) were compared for each phase of the task (Discrimination, Recall, and Reversal). Omission trials did not count towards trials to criterion. In addition, for the Reversal phase we separated errors in which mice chose the odor that was rewarded during Discrimination (Odor 1) into two types: 1) perseverative errors occurred when Odor 1 was chosen prior to the first correct trial and 2) regressive errors occurred when Odor 1 was chosen after the first correct trial during the Reversal phase. To compare the relative proportion of these error types within mice, we calculated Reversal error bias as (perseverative – regressive errors)/(perseverative + regressive errors). Therefore, a value > 0 indicates a bias for perseverative errors whereas a value < 0 indicates a bias for regressive errors. Finally, we examined how quickly mice accumulated rewards after the first correct trial during the Reversal phase by aligning trial histories to the first correct trial and summing rewarded trials across the subsequent 8 trials. Data were fit by linear regression for each group and the slope of the lines compared to determine whether groups significantly differed in their rate of reward accumulation. Behavioral data from 14 of the 16 pOVX females and 10 of the 15 sham females presented here were included in a previously published study examining sex differences of prepubertal gonadectomy on approach-avoidance behaviors, but latent decision variables were not examined (Delevich et al., 2020a).

### Reinforcement learning modeling of 4 choice odor-based reversal task

We modeled Discrimination and Reversal phase behavior using a reinforcement learning model driven by an iterative error-based rule (Rescorla and Wagner, 1972; Sutton and Barto, 2020). On each trial, the model updates the estimated value *V*(*o_i_*, *t*) of the chosen odor *o_i_* according to:

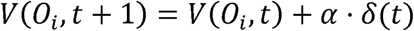

Where the prediction error *δ*(*t*) is the difference between the experience outcome *r*(*t*) and the current estimated value of the chosen odor:

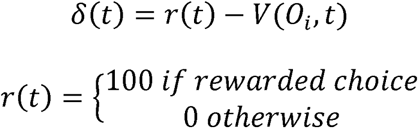

Values of unchosen odors were not updated. The learning rate parameter *α* (0<*α* < 1) scales the prediction error to determine the magnitude of the value update.

Because mice exhibit innate odor preferences, initial odor values *V*(*o_i_*, 0) = *v_i_* werefixed to values estimated from each animal’s choice frequencies during the first four trials of Discrimination (probability of choosing each odor × 100) [38] and were held constant across animals.

Trial-by-trial choice probabilities were computed by passing the odor value estimates through a softmax function:

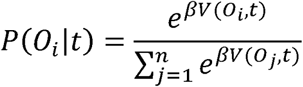

The inverse temperature parameter (β), (referred to throughout the text as the explore/exploit parameter) determines the degree to which choices are guided by current value estimates: high β produces strongly value-dependent (exploitative) choice, whereas low β produces more stochastic (exploratory) choice.

For RL modeling, trial histories from Discrimination and Recall phases were concatenated to create one Discrimination phase trial history. We compared the alternative models using AIC (H. Akaike, 1974) and found that the best fit model included phase-specific (non-zero) α and β parameters; all RL model comparisons for pOVX and sham females are presented in Table S1. To assess model performance, trial-by-trial behavioral data was recovered using the best fit parameters for each animal, and average recovered choices to criterion for Discrimination and Reversal phases (100 simulations/animal) were plotted against the actual choices to criterion for each animal.

### Stereotaxic Virus Injection

Female D2-Cre mice (6–8 weeks) were deeply anesthetized with 5% isoflurane (vol/vol) in oxygen and placed into a stereotactic frame (Kopf Instruments; Tujunga, CA) upon a heating pad. Anesthesia was maintained at 1–2% isoflurane during surgery. An incision was made along the midline of the scalp and small burr holes were drilled over each injection site. Virus was delivered via microinjection using a Nanoject II injector (Drummond Scientific Company; Broomall, PA). Injection coordinates for DMS were (in mm from bregma): 0.90 anterior, +/-1.4 lateral, and -3.0 from surface of the brain. Adeno-associated viruses (AAVs) were produced by Addgene viral service and had titers of >10^12^ genome copies per mL. For chemogenetic manipulations, mice were bilaterally injected with 0.5 uL of rAAV8-hSyn-DIO-mCherry, rAAV8-hSyn-DIO-hM3Dq-mCherry, or rAAV8-hSyn-DIO-hM4Di-mCherry. Mice were given subcutaneous injections of meloxicam (10 mg/kg) during surgery and 24 and 48 hours after surgery. Mice were group-housed before and after surgery and 4–6 weeks were allowed for viral expression before behavioral training or electrophysiology experiments.

### Drugs

Clozapine-N-oxide was generously provided by the NIMH Chemical Synthesis and Drug Supply Program (NIMH C-929). CNO was made fresh each day and dissolved in DMSO (0.5% final concentration) and diluted to 0.1 mg/mL in 0.9% saline USP.

### Electrophysiology

Mice were deeply anesthetized with an overdose of ketamine/xylazine solution and perfused transcardially with ice-cold cutting solution containing (in mM): 110 choline-Cl, 2.5 KCl, 7 MgCl2, 0.5 CaCl2, 25 NaHCO3, 11.6 Na-ascorbate, 3 Na-pyruvate, 1.25 NaH2PO4, and 25 D-glucose, and bubbled in 95% O2/5% CO2. 300 µm thick coronal sections were cut in ice-cold cutting solution before being transferred to ACSF containing (in mM): 120 NaCl, 2.5 KCl, 1.3 MgCl2, 2.5 CaCl2, 26.2 NaHCO3, 1 NaH2PO4 and 11 Glucose. Slices were bubbled with 95% O2/ 5% CO2 in a 37°C bath for 30 min and allowed to recover for 30 min at room temperature before recording. All recordings were made using a Multiclamp 700B amplifier and were not corrected for liquid junction potential. The bath was heated to 32°C for all recordings. Data were digitized at 20 kHz and filtered at 1 or 3 kHz using a Digidata 1440 A system with pClamp 10.2 software (Molecular Devices, Sunnyvale, CA, USA). Only cells with access resistance of <25 MΩ were retained for analysis. Cells were discarded if parameters changed more than 20%. Data were analyzed using pClamp or R (RStudio 0.99.879; R Foundation for Statistical Computing, Vienna, AT).

Whole-cell current clamp recordings were performed using a potassium gluconate-based intracellular solution (in mM): 140 K Gluconate, 5 KCl, 10 HEPES, 0.2 EGTA, 2 MgCl2, 4 MgATP, 0.3 Na2GTP, and 10 Na2-Phosphocreatine. Alexa Fluor 594 (40 µM) was added to the internal solution to enable morphological confirmation of SPN identity following recording. In order to block NMDA and AMPA-mediated currents, 5 µM AP5 and 25 µM NBQX were added to the ACSF, respectively for intrinsic excitability data in Figure 2. For all recordings, cells were allowed to stabilize for 2 min after break in and prior to any current injection. For current clamp recordings to test the effect of CNO in Gq-DREADD- expressing vs. mCherry-expressing D2R(+) neurons, baseline input-output curves were collected before 5 minute wash-on of 10 µM CNO.

### Histology

Mice were transcardially perfused with PBS followed by 4% PFA in PBS. Following 24h postfixation, coronal brain slices (75 µm) were sectioned using a vibratome (VT100S Leica Biosystems; Buffalo Grove, IL). To confirm viral targeting, we performed a standard immunohistochemical procedure using a primary antibody against red fluorescence protein (RFP) (rabbit, Rockland 600-401-379; 1:1000) to enhance the mCherry signal expressed in mice transduced with rAAV8-hSyn-DIO-DREADD-mCherry or rAAV8-hSyn-DIO-mCherry. Sections were counterstained with DAPI (Life Technologies; Carlsbad, CA). Images were acquired with a Zeiss Axio Scan.Z1 epifluorescence microscope (Molecular Imaging Center, UC Berkeley) at 10x magnification and viewed using FIJI (ImageJ). Anatomical regions were identified according to the Mouse Brain in Stereotaxic Coordinates by Franklin and Paxinos and the Allen Institute Mouse Brain Atlas.

### Statistics and Data Analysis

For comparisons between two groups, unpaired two-tailed t-tests were used when data were normally distributed, and Welch’s correction was applied when variances were unequal. Normality was assessed using the D’Agostino & Pearson test. For experiments involving three groups, one-way ANOVA was used for normally distributed data, whereas Kruskal-Wallis tests were used when normality assumptions were violated. Significant omnibus effects were followed by two-tailed Fisher’s LSD or Dunn’s post hoc comparisons, respectively. Two-way RM ANOVA was used when experiments included both within- and between-subject factors (e.g. task phase × treatment), followed by planned two-tailed Fisher’s LSD post hoc comparison. Post hoc comparisons were not corrected for multiple comparisons, because the number of planned comparisons was limited and hypothesis driven. Effect sizes were reported as Cohen’s *d* for pairwise comparisons, eta squared (η^2^) or partial eta squared (η_p_^2^) for ANOVA effects, and epsilon squared (ε^2^) for Kruskal-Wallis tests. Rank-based post hoc comparisons are reported with effect size *r*. Throughout the paper, p=0.05 was used as the criterion for a significant statistical difference unless noted otherwise. Data are expressed as mean ± SEM unless noted otherwise.

## Results

### Prepubertal ovariectomy increases exploratory choice during reversal learning

We performed sham surgery or pOVX on female C57BL/6 mice at postnatal day 25 (P25), prior to puberty onset, and trained them in an odor-based reversal task between P60-70 (Fig. 1A). The odor-based reversal task consisted of two main phases: 1) a Discrimination phase during which mice learned through trial and error that one of four scented pots of wood shavings contained a buried food reward and 2) a Reversal phase in which the odor-reward contingency was reversed (Fig. 1B). Sham females were not staged for estrous cycle, and both groups performed similarly in the Recall phase (Supplementary Fig. 1). When comparing Discrimination and Reversal, there was a significant effect of task phase but not treatment on trials to reach criterion [task phase: F(1,29)= 6.31, p= 0.018, η_p_^2^= 0.18; treatment: F(1,29)= 0.11, p= 0.74, η_p_^2^= 0.005; task phase × treatment: F(1,29)= 0.05, p= 0.83, , η_p_^2^= 0.002] (Fig. 1C).

**Figure 1.**
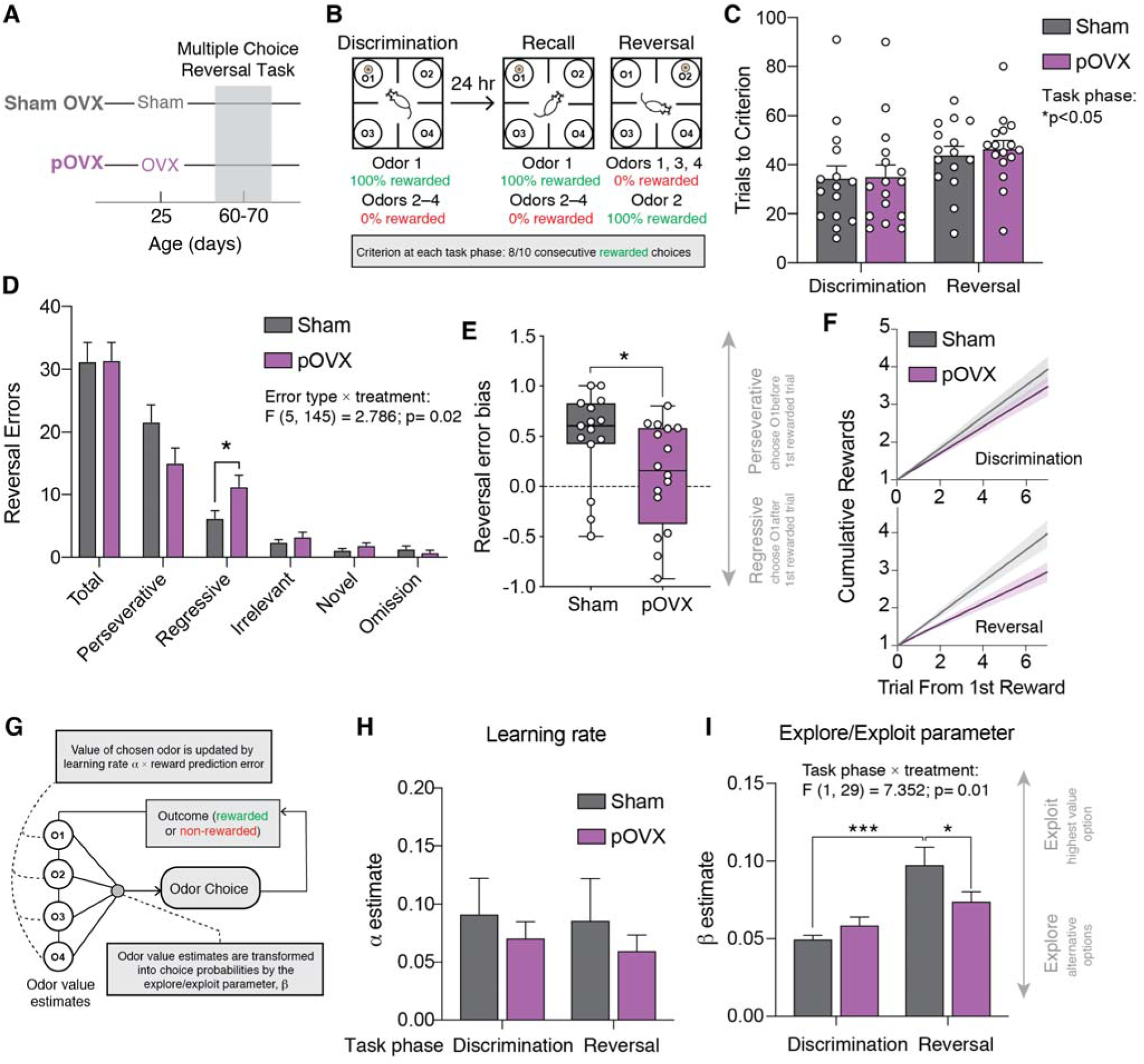
Prepubertal OVX is associated with more exploratory reversal learning strategy in female mice. **(A)** Female C57BL/6 mice underwent OVX or sham surgery at P25 and were trained in the multiple choice reversal task in adulthood (P60-70). **(B)** Mice were trained to a criterion of 8/10 correct consecutive choices to Odor 1 during Discrimination. 24 hours later they were tested for Recall of the previous day’s rule before immediately advancing to a Reversal phase during which Odor 2, rather than Odor 1, was rewarded. Reversal criterion was reached when mice made 8/10 correct consecutive choices to Odor 2. **(C)** There was a main effect of task phase on trials to criterion but no effect of treatment (main effect of task phase F(1, 29) = 6.30, p<0.05, Two-way RM ANOVA). **(D)** There was a significant effect of treatment and error type on the number of reversal errors (treatment × error type interaction: F(5, 145) = 2.79, p<0.05, Two-way RM ANOVA). pOVX females made significantly more regressive errors compared to sham females (11.25 ± 1.8 vs. 6.13 ± 1.2, p<0.05, uncorrected Fisher’s LSD). **(E)** pOVX females had a significantly lower Reversal error bias (perseverative – regressive errors)/(perseverative + regressive errors) compared to sham females (0.11 ± 0.13 vs. 0.49 ± 0.12, p<0.05, unpaired t-test). **(F)** Sham females accumulated rewards after the first correct Reversal trial faster than pOVX females (best fit line with 95% C.I. plotted). **(G)** RL model applied to odor-based multiple choice reversal task. Schematic based on (Verharen et al., 2019b). **(H)** Best-fit α learning rate estimates did not significantly differ by task phase or treatment. **(I)** There was a significant interaction between task phase and treatment group on the best-fit explore/exploit β parameter ( task phase × treatment interaction: F(1,29)= 7.35, p<0.05, Two-way ANOVA). Post hoc comparisons revealed that β parameter estimates were significantly higher during Reversal compared to the Discrimination phase for sham (p<0.0001, Sidak’s multiple comparisons test) but not pOVX females. In addition, Reversal phase β parameter estimates were significantly lower in pOVX females compared to sham (p<0.05, Sidak’s multiple comparisons test).

Next, we more closely examined the types of errors that mice made during Reversal. Error types included those made to the previously rewarded odor, which we divided into 2 subtypes: perseverative (errors made before the first correct trial) and regressive (errors made after first correct trial). Perseverative errors reflect a tendency to stick to a previously learned rule, whereas regressive errors reflect a failure to acquire or maintain the new rule. There was a significant interaction between error type and treatment group [F(5,145)=2.79, p=0.02, η_p_^2^= 0.088] (Fig. 1D). Post hoc analyses revealed that pOVX females made significantly more regressive errors compared to sham females (p=0.03, d=0.83, uncorrected Fisher’s LSD). We next examined the pattern of perseverative and regressive errors made by individual mice. Sham females exhibited a significantly higher ratio of perseverative to regressive errors (Reversal error bias) compared to pOVX females (sham vs. pOVX females: t(29)=2.12, p= 0.04, d=0.76) (Figure 1E). Finally, we observed that sham females accumulated rewards at a significantly higher rate after the first rewarded trial compared to pOVX females during Reversal but not Discrimination (Figure 1F). These data suggest that sham females and pOVX females reach criterion in the reversal task using different trial-by-trial strategies.

We next turned to computational modeling to assess if differences observed in the Reversal phase between sham and pOVX females arise from a difference in odor value updating, a difference in choice policy, or a combination of both. To do so, we fit trial-by-trial behavioral data with an RL model and used the maximum log likelihood to determine the parameters that best fit each animal’s behavior. The best fit model included phase-specific parameters for the learning rate α and the explore/exploit inverse temperature parameter β (Fig. 1G) (see Supplementary Table 1 for alternate model comparison). There were no significant effects of task phase or treatment on the learning rate parameter α [task phase × treatment: F(1,29)=0.013, p=0.91, η_p_^2^<.001] (Fig. 1H). By contrast, there was a significant interaction between task phase and treatment for the explore/exploit parameter β [task phase × treatment: F(1,29) = 7.35, p= 0.011, η_p_^2^=.202]. In sham but not pOVX female mice, the explore/exploit parameter was significantly higher during the Reversal phase compared to Discrimination phase (sham Reversal vs. Discrimination: p<0.0001, d=2.04; pOVX Reversal vs. Discrimination p= 0.07, d=0.66, uncorrected Fisher’s LSD) and Reversal phase explore/exploit parameter was significantly lower in pOVX vs. sham females (pOVX vs. sham: p= 0.022, d=0.85, uncorrected Fisher’s LSD) consistent with pOVX females employing a more exploratory choice policy compared to sham females (Fig. 1I).

### Prepubertal OVX is associated with increased intrinsic excitability of D2R(+) SPNs

The DMS has been implicated in action selection and determining choice policy, and previous studies have found evidence that estrous cycle modulates the intrinsic excitability of striatal SPNs. Furthermore, several lines of evidence suggest that D2R(+) iSPNs: 1) are modulated by ovarian hormones (Krentzel et al., 2021; Le Saux et al., 2006; Le Saux and Di Paolo, 2005) and 2) can influence explore/exploit balance during decision making (Delevich et al., 2022; Kwak et al., 2014; Lee et al., 2015). We therefore investigated whether changes in the intrinsic excitability of D2R(+) SPNs within DMS may contribute to sham vs. pOVX differences in choice policy during reversal learning. We performed whole-cell current clamp recordings of visually identified eGFP+ and neurons within the DMS of adult D2-eGFP transgenic female mice who underwent pOVX or sham surgery and unmanipulated female mice in the presence of the excitatory synaptic blockers NBQX and AP5 (Fig. 2A-C). AlexaFluor-594 was included in the internal solution, and all cells included in analysis were confirmed to have spinous morphology. We found a main effect of treatment on D2-eGFP(+) SPN input resistance [main effect of treatment: H= 8.76, p= 0.0125, ε^2^=.19] with pOVX females exhibiting higher input resistance compared to sham and unmanipulated females (p<0.05, *r*=.47, Dunn’s multiple comparisons test) (Fig. 2D). When we injected a series of positive current steps (Fig. 2E), we found that the minimum current required to trigger an action potential (rheobase) differed significantly across groups [main effect of treatment: F(2,35)= 7.97, p=0.0014, η^2^=.31, Two-way RM ANOVA], with pOVX females exhibiting lower rheobase compared to sham-operated (p=0.003, d=1.37, Sidak’s multiple comparisons test) and unmanipulated mice (p=0.007, d=1.39, Sidak’s multiple comparisons test) (Fig. 2F). In addition, there was a significant interaction between treatment and current on spike output [F(98, 1715)= 2.517, p<0.0001, η_p_^2^=.13, Two-way RM ANOVA] (Fig. 2G). While input-output curves were shifted leftward in pOVX compared to sham and unmanipulated females, there was no significant effect of treatment on maximum firing rate [F(2, 18.68)= 0.10, p= 0.90] (Fig. 2H). Finally, resting membrane potential (RMP) did not differ across treatments [F(2, 35)= 2.172, p= 0.129, η2=.11] (Fig. 2I). These data indicate that D2R(+) SPNs within the DMS are more intrinsically excitable in pOVX females compared to unstaged sham and unmanipulated female mice.

**Figure 2.**
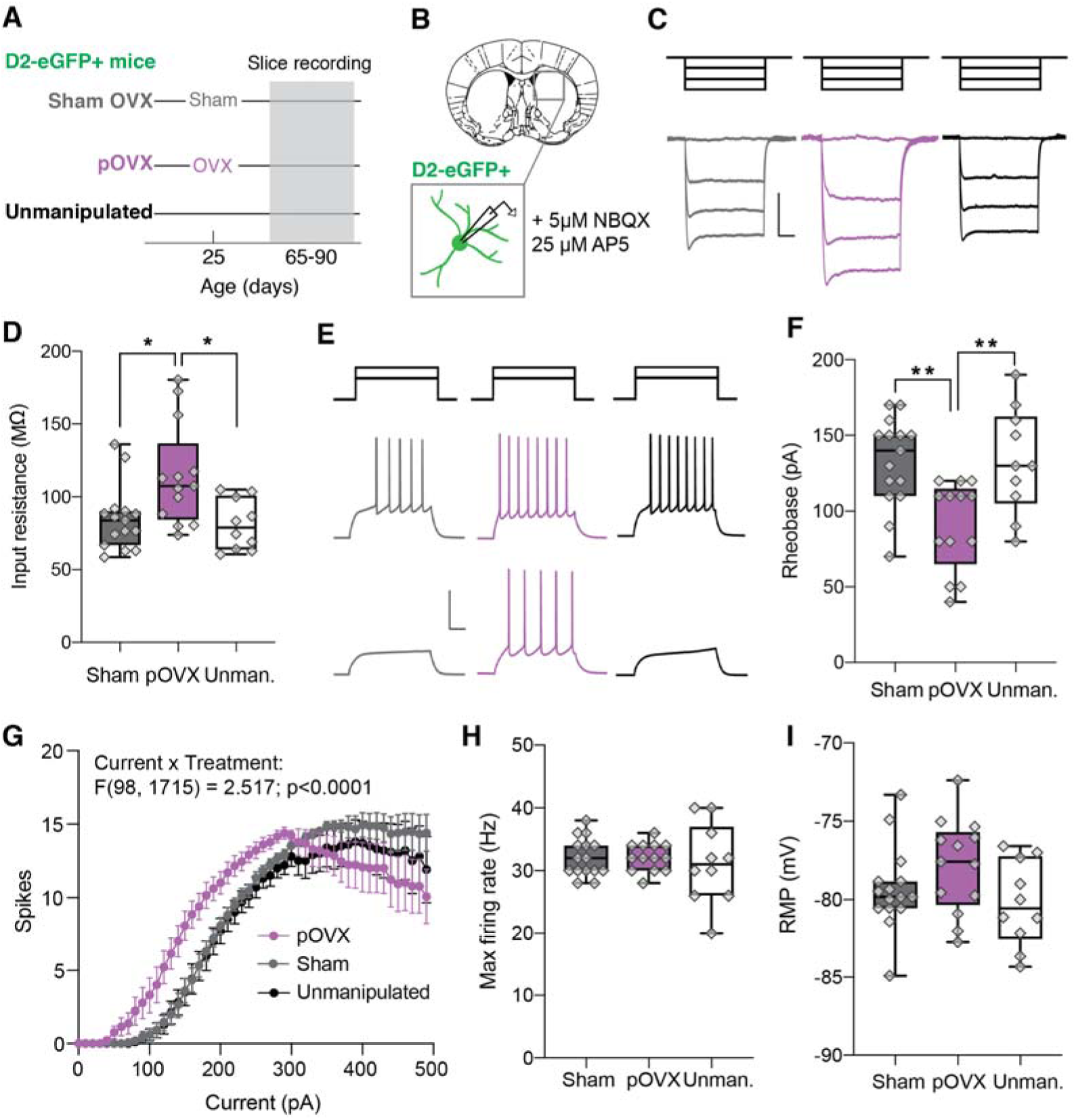
Prepubertal OVX increases D2R(+) SPN intrinsic excitability in the presence of synaptic blockers. **(A)** At P25 D2-eGFP(+) female mice underwent sham or pOVX surgery, while a third group of female D2-eGFP(+) mice received no surgery. **(B)** Whole-cell current clamp recordings were made from visually identified D2-eGFP(+) SPNs within the DMS from all groups in adulthood (P65–90). **(C)** Representative responses to negative current steps (-150, - 100, -50, 0 pA) in D2R(+) SPNs from sham, pOVX, and unmanipulated females. Scale bar: 100 ms, 5 mV. **(D)** D2-eGFP(+) SPNs in pOVX female mice had higher input resistance compared to sham and unmanipulated females. **(E)** Representative responses to positive current steps (120, 180 pA) in D2R(+) SPNs from sham, pOVX, and unmanipulated females. Scale bar: 100 ms, 50 mV. **(F)** Decreased rheobase of D2-eGFP(+) SPNs was observed in pOVX compared to sham and unmanipulated females. **(G)** Spike number across sequential depolarizing current steps (10–500 pA) for D2-eGFP(+) SPNs. Increased spiking was observed in pOVX compared to sham and unmanipulated females (current x treatment interaction: F(98, 1715)= 2.52, p<0.0001, Two-way RM ANOVA). **(H)** No difference in maximum firing rate was observed across treatment groups. **(I)** No difference in RMP was observed across treatment groups. *p<0.05, **p<0.01; n/N = 15/5, 13/5, and 10/3 for sham, pOVX, and unmanipulated mice, respectively.

### Chemogenetic activation of D2R(+) SPNs in DMS reduces perseverative bias and promotes a more exploratory reversal strategy in female mice

Given that OVX females exhibit a more exploratory reversal strategy and greater intrinsic excitability of D2R(+) SPNs in DMS, we next asked whether experimentally increasing D2R(+) SPN intrinsic excitability would similarly bias intact female mice towards increased exploration during the Reversal phase. Female D2-Cre mice were bilaterally infused with 0.5 µL of Cre-dependent DREADD virus (hM4Di-mCherry or hM3Dq-mCherry) and trained 4-6 weeks later in the 4 choice odor-based reversal learning task (Fig. 3A). Female mice expressing Cre-inducible mCherry were used to control for any effects of surgery, AAV infection, and clozapine-N-oxide (CNO) administration on behavior. To examine how CNO activation of hM3Dq expressed by D2R(+) SPNs in DMS alters their activity, we performed whole-cell current clamp recordings of identified mCherry+ neurons in mice that expressed the excitatory DREADD hM3Dq or mCherry alone (Fig. 3B). Briefly, spike output in response to depolarizing steps (0–360 pA, 20 pA steps) was recorded from visually identified mCherry+ neurons in D2-mCherry or D2-hM3Dq-mCherry expressing SPNs in DMS (Fig. 3A-D). Next, 10 µM CNO was bath-applied for 5 minutes and spike output to the same sequential series of depolarizing current steps was recorded (Fig. 3A-D). There was no significant interaction between current step and drug on spike output in D2-mCherry expressing SPNs [current x drug: F(18,36)=0.89, p=0.59, η_p_^2^=.309, Two-way RM ANOVA] (Fig. 3C) but there was a significant interaction between current and drug on spike output in D2-hM3Dq-mCherry SPNs [current x drug: F(18,36)=3.93, p=0.0002, η_p_^2^=.98, Two-way RM ANOVA] (Fig. 3D). Finally, there was a significant interaction between virus and drug on rheobase [F(1,4)= 16.0, p=0.016, η_p_^2^=.80, Two-way RM ANOVA] (Fig. 3E).

**Figure 3.**
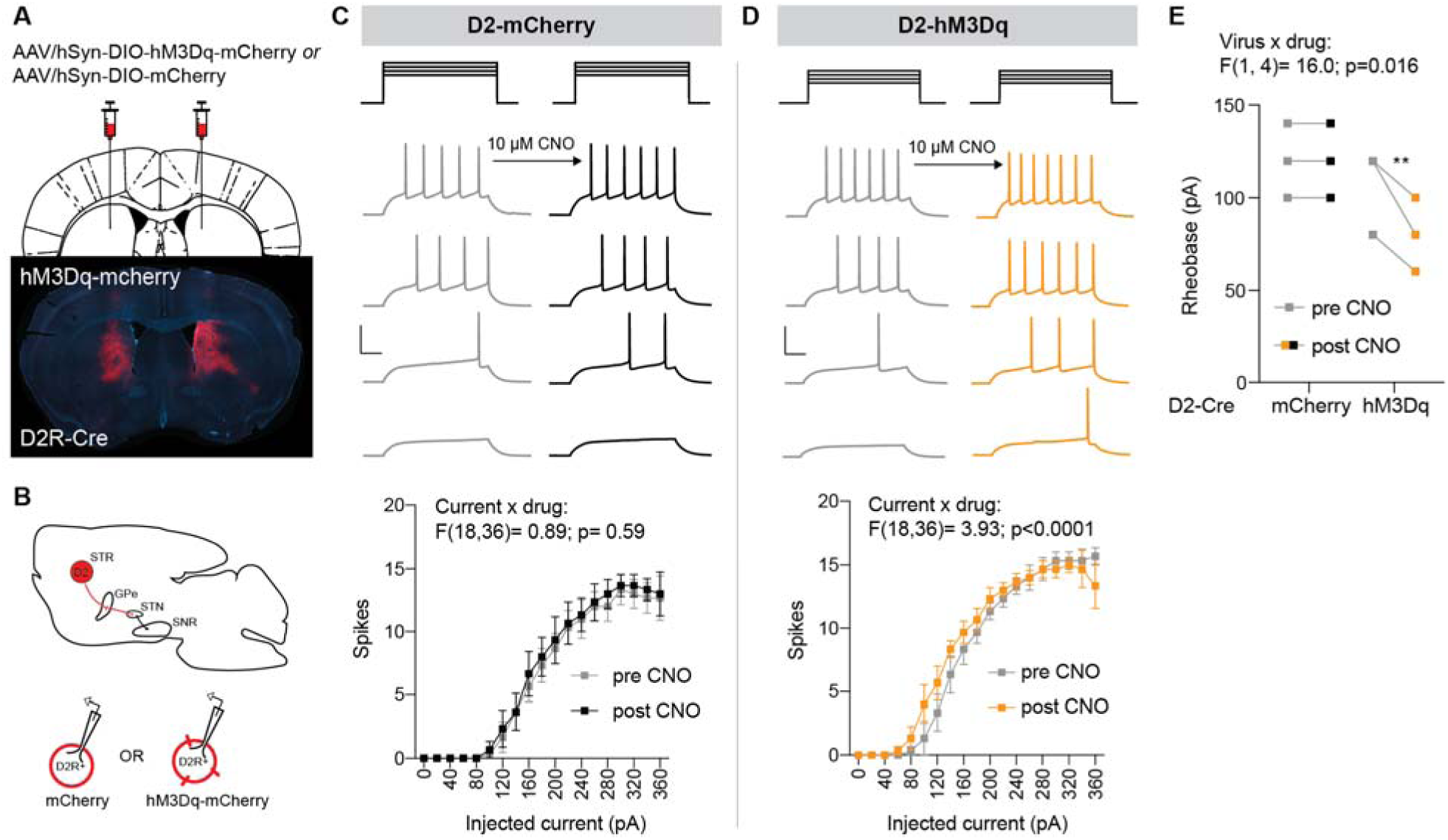
CNO increases intrinsic excitability of hM3Dq-expressing D2R(+) iSPNs. **(A)** Schematic of injection and representative brain section showing hM3Dq-mCherry expression in the DMS of D2-Cre mouse. **(B)** Schematic of indirect pathway expression (sagittal view) and whole-cell patch-clamp configuration of mCherry+ or hM3Dq-mCherry+ iSPNs in female D2-Cre mice. **(C)** Top panel: representative responses to positive current steps (100, 120, 140, 160 pA) in mCherry+ iSPNs before and after CNO wash on. Scale bar: 100 ms, 50 mV. Bottom panel: no significant interaction between current step and CNO treatment on spike output in D2R(+) mCherry-expressing iSPNs. **(D)** Top panel: representative responses to positive current steps (100, 120, 140, 160 pA) in hM3Dq-mCherry+ iSPNs before and after CNO wash on. Scale bar: 100 ms, 50 mV. Bottom panel: significant interaction between current step and CNO treatment on spike output in D2R(+) mCherry-expressing iSPNs (current step × drug, p<0.0001, Two-way RM ANOVA). **(E)** Summary of CNO wash on effect on rheobase (virus × drug, p<0.05, Two-way RM ANOVA).

Prior to Discrimination training all mice received i.p. injections of saline (Fig. 4A) and learned through trial and error that one of four presented odors indicated the location of a buried food reward. Mice completed the Discrimination task phase when they selected the rewarded odor (Odor 1) on 8/10 consecutive trials. Twenty-four hours later, all groups were administered CNO (1.0 mg/kg, i.p.) and tested for their recall of discrimination learning followed immediately by a Reversal phase in which Odor 1 was no longer rewarded and Odor 2 became rewarded. There was a significant effect of task phase on trials to criterion [Reversal vs. Discrimination; F(1,17)= 16.58, p= 0.0008, η_p_^2^=.494, Two-way RM ANOVA] but no significant effect of virus [F(2,17) = 0.37, p= 0.69, η_p_^2^=.03, Two-way RM ANOVA] or interaction between virus and task phase [F(2,17)= 0.09, p=0.92, η_p_^2^=.01, Two-way RM ANOVA] (Fig. 4B). While there was no significant effect of chemogenetic manipulation on Reversal phase trials to criterion, we found a significant interaction between virus and error type during Reversal [F(10,85)= 2.721, p= 0.006, η_p_^2^=.24, Two-way RM ANOVA] (Fig. 4C) that was absent during Discrimination when mice were on saline (Supplementary Figure 2). D2-hM3Dq mice made significantly fewer perseverative errors compared to D2-mCherry (p= 0.015, d=1.55, uncorrected Fisher’s LSD) and D2-hM4Di groups (p= 0.028, d=1.88, uncorrected Fisher’s LSD) and made significantly more regressive errors compared to D2-hM4Di mice (p= 0.03, d=1.72, uncorrected Fisher’s LSD) (Fig. 4C). We next examined whether chemogenetic manipulation of D2R(+) neurons in the DMS altered Reversal error bias within mice. There was a significant effect of virus on Reversal error bias (H= 9.06, p= 0.005, ε^2^=.42, Kruskal-Wallis test), with D2-hM3Dq mice having a significantly lower Reversal error bias compared to D2-mCherry (p=0.019, *r*=.61 uncorrected Dunn’s test) and D2-hM4Di groups (p= 0.005, *r*=.85, uncorrected Dunn’s test) (Fig. 4D), consistent with a greater tendency to make regressive errors compared to perseverative errors. These data suggest that chemogenetic activation of D2R(+) neurons in the DMS produced a pattern of reversal phase choice behavior that was similar to the effect seen in pOVX mice.

**Figure 4.**
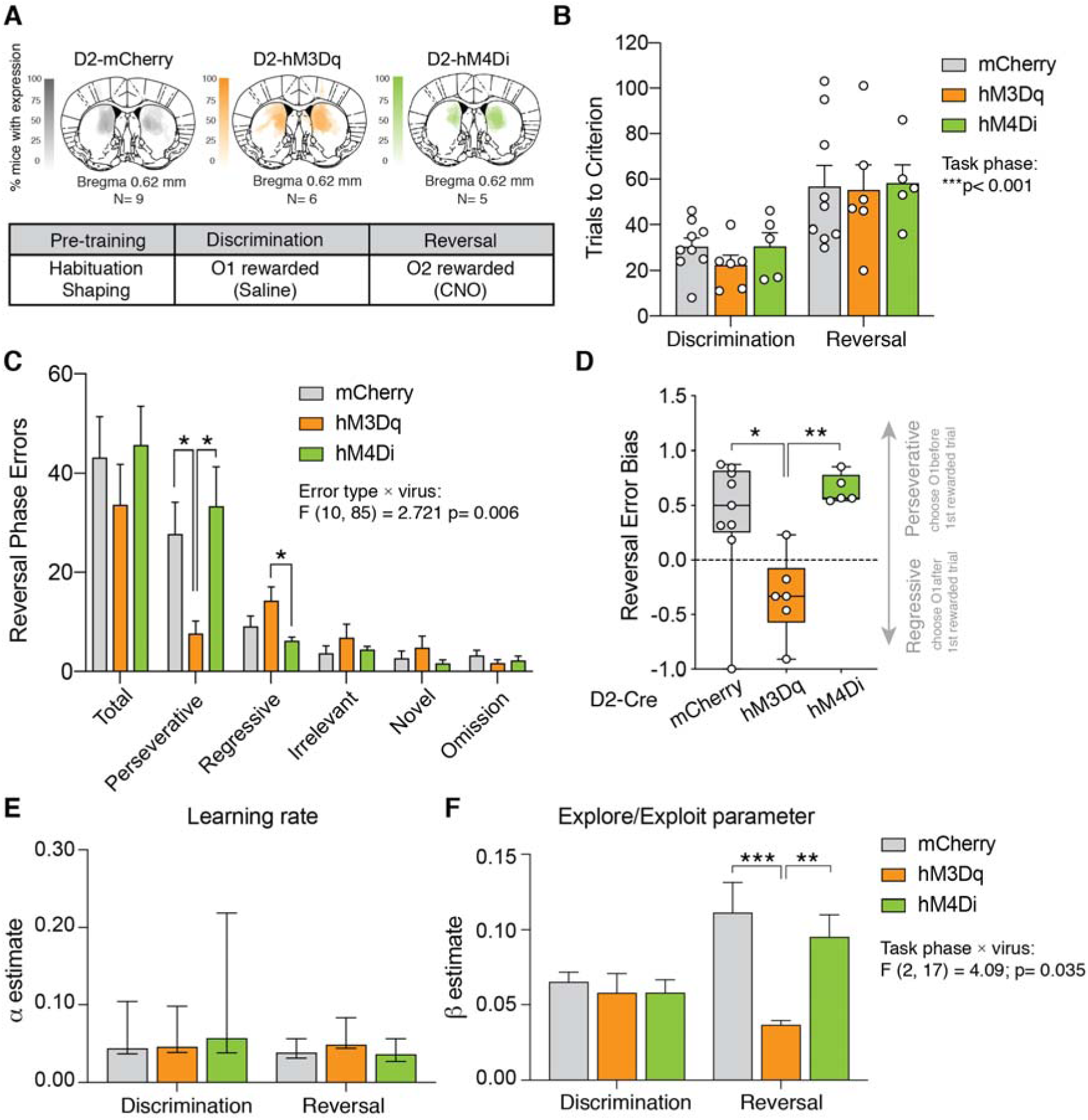
Chemogenetic activation of D2R(+) neurons in DMS promotes a more exploratory reversal strategy in intact female mice. **(A)** Top panel: schematic illustrating injection site and viral spread female D2-Cre DIO-mCherry (N=9), DIO-hM3Dq (N=6), and DIO-hM4Di (N=5) mice. Bottom panel: summary of behavioral training. **(B)** There was a main effect of task phase but no effect of virus on trials to criterion (main effect of task phase: F(1,17)= 16.58, p<0.001, Two-way RM ANOVA). **(C)** There was a significant interaction between error type and virus on reversal errors (error type × manipulation interaction: F(10,85)= 2.72, p<0.01, Two-way RM ANOVA) with D2-hM3Dq mice making fewer perseverative errors (p<0.05, uncorrected Fisher’s LSD) compared to D2-mCherry and D2-hM4Di mice and more regressive errors compared to D2-hM4Di mice (p<0.05, uncorrected Fisher’s LSD). **(D)** There was a main effect of virus on Reversal error bias (H=9.06, p<0.01 Kruskal-Wallis test), with D2-hM3Dq mice showing reduced bias for perseverative errors compared to D2-mCherry (p<0.05, uncorrected Dunn’s test) and D2-hM4Di mice (p<0.01, uncorrected Dunn’s test). **(E)** Best-fit α learning rate did not significantly differ by task phase or virus. **(F)** There was a significant interaction between task phase and treatment group on the best-fit explore/exploit parameter β (Two-way ANOVA task phase × treatment interaction: F(2,17)= 4.09, p<0.05). Post hoc comparisons revealed that β parameter estimates were significantly higher during Reversal compared to Discrimination phase for D2-mCherry mice (p<0.01, uncorrected Fisher’s LSD) but not D2-hM3Dq mice (p=0.28, uncorrected Fisher’s LSD). In addition, Reversal phase β parameter estimates were significantly lower in D2-hM3Dq mice compared to D2-mCherry and D2-hM4Di mice. *p<0.05, **p<0.01, ***p<0.001.

Finally, we applied RL modeling to determine whether similar changes in decision-making parameters might explain the pattern of reversal behavior we observed in pOVX female mice and D2-hM3Dq female mice. Fitting the same RL model (task phase-specific α and β parameters; see Methods), a Two-way RM ANOVA revealed no significant interaction between virus and task phase on learning rate α [F(2,17)= 0.49, p=0.62, η_p_^2^=.054] (Fig. 4E), but a significant interaction between virus and task phase for the explore/exploit parameter β [F(2,17)= 4.09, p=0.035 η_p_^2^=.33] (Fig. 4F). The Reversal phase explore/exploit parameter was significantly lower in D2-hM3Dq mice compared to D2-mCherry (p=0.0003, d=2.12, uncorrected Fisher’s LSD) and D2-hM4Di (p= 0.0095, d=1.66, uncorrected Fisher’s LSD) (Fig. 4F). These data suggest that chemogenetic activation of D2R(+) neurons within DMS biases choice strategy in female mice to be more exploratory during reversal learning. Moreover, chemogenetic activation of D2R(+) neurons within the DMS produced behavior in female mice that mimicked the behavioral pattern seen in OVX females, including a reduction in Reversal error bias during reversal learning and a reduced explore/exploit β parameter consistent with a less exploitative, more exploratory choice policy. Taken together with evidence that D2R(+) iSPNs within the DMS are more intrinsically excitable in pOVX compared to sham females, these data support a model whereby pOVX biases reversal learning strategy towards exploration by modulating iSPN intrinsic excitability within DMS.

## Discussion

We found that pOVX altered how female mice solved a reversal learning task. Using an RL model fit to trial-by-trial behavioral data, we found that intact females flexibly adjusted their choice policy as the task progressed: they employed a relatively exploratory policy during initial discrimination learning and shifted toward a more exploitative policy during Reversal, captured by a phase-dependent increase in the inverse temperature parameter (β). By contrast, pOVX females maintained a comparatively exploratory choice policy across both phases. This absence of task-phase-dependent modulation of choice policy was accompanied by increased intrinsic excitability of D2R(+) iSPNs in the DMS, a region that is implicated in regulating action selection and choice policy. We then sought to mimic this effect using chemogenetics. We demonstrated that chemogenetic activation of D2R(+) neurons *in vitro* similarly enhanced iSPN intrinsic excitability in slices from female brains. In addition, activation of DMS D2R(+) neurons *in vivo* decreased the ratio of perseverative to regressive errors and prevented the typical shift toward a more exploitative choice policy in the Reversal phase, captured by a flatter β profile across task phases. Together, these data suggest that two distinct manipulations—pOVX and hM3Dq-mediated activation of DMS D2R(+) neurons—converge on a shared physiological phenotype of enhanced DMS iSPN intrinsic excitability that is associated with the maintenance of a more exploratory choice strategy.

Our data are consistent with studies that manipulate D2Rs and model choice policy. Germline D2R knockout (Kwak et al., 2014), systemic D2R antagonist administration (Eisenegger et al., 2014), and intrastriatal D2R antagonist infusion (Lee et al., 2015) are each associated with less value-dependent, more exploratory choice policy. However, none of these studies could rule out the contribution of presynaptic D2 autoreceptors, which is important given the apparent role of tonic dopamine in modulating explore/exploit balance (Beeler et al., 2010; Cinotti et al., 2019; Humphries et al., 2012), but see (Costa et al., 2014). Our chemogenetic manipulation experiments (which do not infect D2R(+) dopamine axon terminals) clearly demonstrate that activation of D2R(+) neurons within DMS is sufficient to bias performance towards exploration.

We speculate that there are two likely circuit mechanisms downstream of D2R(+) iSPNs that may be responsible for promoting exploratory choice policy. The first involves local lateral connections from iSPNs to direct pathway SPNs (dSPNs) and the second involves the interface of the direct and indirect pathways in basal ganglia output centers such as the substantia nigra pars reticulata (SNr). One recent study showed that systemic injection of the D2R antagonist raclopride induced dopamine-dependent transcriptional activation in iSPNs that opposed the activation of dSPNs, suggesting that iSPN-to-dSPN transmodulation contributes to behavioral flexibility (Matamales et al., 2020). Therefore, elevated iSPN activity induced either by pOVX or chemogenetic activation may promote exploratory choice by dampening activity within dSPN ensembles that normally promote the selection of the highest estimated-value option. Opponent interactions between the direct and indirect pathways at convergent downstream targets have similarly been proposed to regulate choice policy (Collins and Frank, 2014).

Studies have shown that the intrinsic properties of striatal SPNs differ between females and males before puberty (Dorris et al., 2015) and fluctuate across the estrous cycle after puberty (Proaño et al., 2018; Willett et al., 2020). Estrous cycle-dependent modulation of dorsal striatal SPN physiology appears to be multifaceted and property-specific rather than reflecting a simple unidirectional change in overall excitability (Willett et al., 2020) In the accumbens, Proaño et al. found that SPN intrinsic excitability was significantly higher during diestrus/metestrus, when circulating estradiol and progesterone levels are low, compared to proestrus/estrus (Proaño et al., 2018) and that estradiol replacement in OVX rats at levels that mimic proestrus/estrus significantly decreased accumbal SPN intrinsic excitability (Proaño and Meitzen, 2020). Together, these studies demonstrate that ovarian hormones dynamically regulate intrinsic electrophysiological properties of striatal SPNs in adulthood across both dorsal and ventral striatal regions. Our findings extend this literature by demonstrating that OVX prior to puberty onset is associated with long-lasting enhancement of intrinsic excitability in D2R(+) SPNs within the DMS. One possible mechanism is through direct estrogen receptor signaling within SPNs, as SPNs express membrane-associated estrogen receptors and estradiol can rapidly modulate SPN intrinsic excitability and excitatory synaptic transmission (Krentzel et al., 2021, 2019; Proaño and Meitzen, 2020). In the dorsal striatum, membrane-associated ERα, ERβ, and GPER-1 receptors have been proposed to regulate SPN physiology through coupling with mGluR signaling pathways (Willett et al., 2020). Alternatively, ovarian hormones may indirectly regulate SPN physiology through modulation of striatal dopamine signaling. Estradiol has also been shown to influence dopamine release and reuptake in the striatum (Calipari et al., 2017; Golden et al., 2025), including the dorsal striatum (Becker, 1990; Becker and Beer, 1986), and dopamine signaling contributes to the postnatal maturation of SPN intrinsic excitability (Lieberman et al., 2018). A recent study found that prepubertal gonadectomy in male wild-derived mice significantly reduced the density of dopamine release sites in the dorsal striatum (Jackson et al., 2025), although downstream effects on SPN physiology were not explored. Together, these findings raise the possibility that pubertal ovarian hormones contribute to the developmental maturation of striatal circuitry through both direct hormonal signaling within SPNs and indirect modulation of dopamine system maturation or function (Lin et al., 2020).

There are several lines of evidence that suggest that ovarian hormones preferentially modulate the activity of D2R(+) iSPNs versus D1R(+) dSPNs. OVX decreases D2 receptor binding in the striatum, and estradiol or treatment with an ERβ agonist counteracts the effect of OVX (Le Saux et al., 2006). Importantly, these manipulations did not alter D2R mRNA expression, suggesting that estradiol modulates D2R binding through a mechanism other than transcriptional regulation of D2R expression. Furthermore, OVX reduces the expression of preproenkephalin, which produces the endogenous opioid peptide enkephalin expressed by iSPNs (Le Saux and Di Paolo, 2005). Again, the effects of OVX on preproenkephalin expression can be reversed by estradiol administration. Together, these findings suggest that ovarian hormones may preferentially influence indirect pathway signaling within the striatum.

At the cellular level, several mechanisms could contribute to the enhanced excitability of D2R(+) SPNs observed following pOVX. Ovarian hormones have been shown to regulate dendritic complexity and spine density in cell types in other brain regions (Chen et al., 2009; Gould et al., 1990; Wallace et al., 2006; Woolley et al., 1990; Ye et al., 2019). The dendrites of D2R(+) SPNs are enriched in Kir2 family inward rectifying K+ channels (Mermelstein et al., 1998; Nisenbaum and Wilson, 1995; Shen et al., 2007; Uchimura et al., 1989), and a reduction in dendritic length/complexity has been associated with reduced Kir2 expression and increased intrinsic excitability (Cazorla et al., 2012; Sebel et al., 2017). Therefore, it would be informative to compare iSPN Kir2 channel currents and dendritic morphology in sham vs. pOVX females. Finally, it is possible that the increase in intrinsic excitability of the D2R(+) SPNs in pOVX females could represent a homeostatic plasticity mechanism that accompanies a reduction in excitatory synaptic inputs to them, but we did not measure synaptic inputs onto D2R(+) SPNs in this study.

There are several limitations to our current study that should be noted. First, we cannot assume that the changes we observed in SPN intrinsic excitability are specific to the dorsomedial region of the striatum or to the D2R(+) SPN cell type. Second, our evidence linking the increased excitability of D2R(+) SPNs in DMS to the loss of task-phase-dependent modulation of choice policy in pOVX females is correlational. In independent experiments we observed that 1) pOVX females did not shift toward a more exploitative choice policy in the Reversal phase of the experiment as was observed in sham adult females, 2) pOVX was associated with elevated intrinsic excitability of D2R(+) SPNs in DMS and 3) that chemogenetic activation of D2R(+) SPNs in DMS in intact females similarly disrupted the task-phase-dependent shift toward a more exploitative choice policy during Reversal. In the future, more direct evidence could be gained by performing manipulation experiments to reduce the activity of DMS D2R(+) SPNs in OVX females, or by recording the activity of these same neurons in pOVX and sham females during behavior. Finally, while we performed OVX prior to puberty onset, we do not know whether the timing of OVX plays an important role in the observed effect on behavior and physiology. We also do not know if and when hormone replacement may rescue the effects of pOVX. Future studies using temporally-targeted hormone replacement strategies will be important for distinguishing potential organizational effects of pubertal ovarian hormone exposure from acute activational effects in adulthood. Still, in light of these limitations, our data suggest future lines of inquiry into the relationship between puberty, ovarian hormones, SPN physiology, and choice policy in value-based decision making.

There is a growing interest in understanding the mechanisms that underlie sex differences in value-based decision making (Cox et al., 2023; Golden et al., 2025; Grissom and Reyes, 2019). A recent study showed that compared to males, female mice employed a more consistent strategy while learning a two-dimensional decision-making task (Chen et al., 2021). This tendency to commit to a strategy once reward contingencies are identified aligns with the shift toward exploitation we observed in sham females during Reversal. Conversely, we observed that pOVX females failed to make this shift, instead maintaining a more exploratory pattern of choice, sticking less to the previously rewarded odor choice and committing more regressive errors than sham females. These findings suggest that ovarian hormones contribute to the female-biased choice strategies utilized during value-based decision making. While previous studies have separately provided evidence that ovarian hormones regulate the intrinsic excitability of SPNs and aspects of value-based decision making, we show for the first time that pOVX disrupts the task-phase-dependent modulation of the explore/exploit balance of choice strategy while also increasing the intrinsic excitability of D2R(+) SPNs in the DMS. These data suggest that pubertal status may influence the flexible adjustment of explore/exploit balance via the modulation of SPN intrinsic excitability within the DMS and highlight a role for ovarian hormones in establishing sex-specific decision-making strategies in adulthood. These findings can inform the basic science of decision making and the study of the many psychiatric disorders that emerge after puberty and exhibit sex differences in their prevalence or manifestation.

## Acknowledgments

We thank Yuting Zhang, Kenechukwu Okwuosa, and Weihang Chen for technical assistance with mouse behavior testing. We thank Dr. Helen Bateup for feedback on the manuscript and Dr. Anne Collins and Wilbrecht lab members for helpful discussion.

## Data Availability Statement

The data supporting the findings of this study are openly available at Figshare: https://doi.org/10.6084/m9.figshare.14783628.v1

