## Supplementary Materials for "Prepubertal ovariectomy alters dorsomedial striatum indirect pathway neuron excitability and explore/exploit balance in female mice"

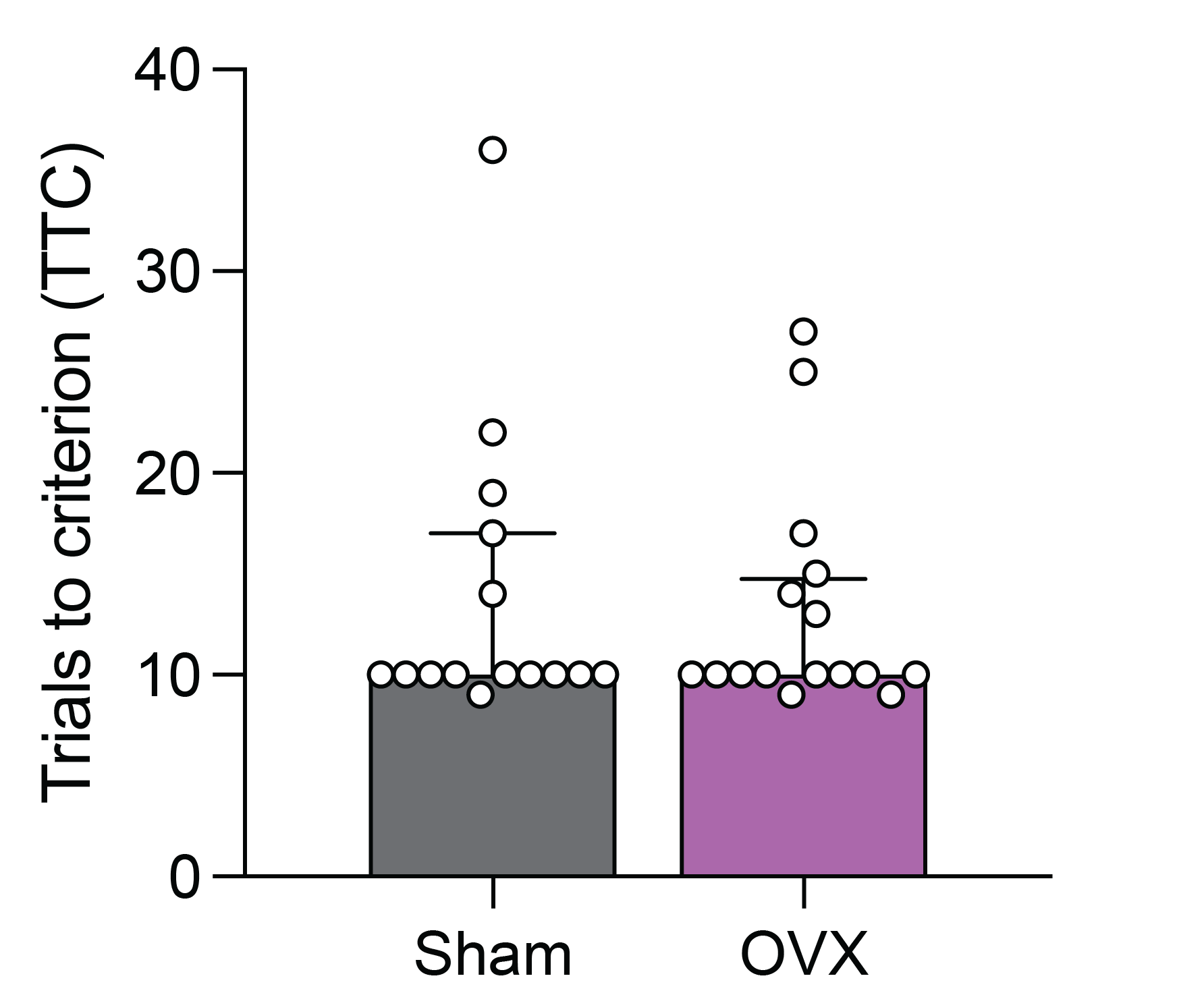

**Supplementary Figure 1.** OVX and sham females do not differ in TTC during the recall phase of the odor-based multiple choice foraging task. p= 0.87, Mann-Whitney U test; N=15,16.

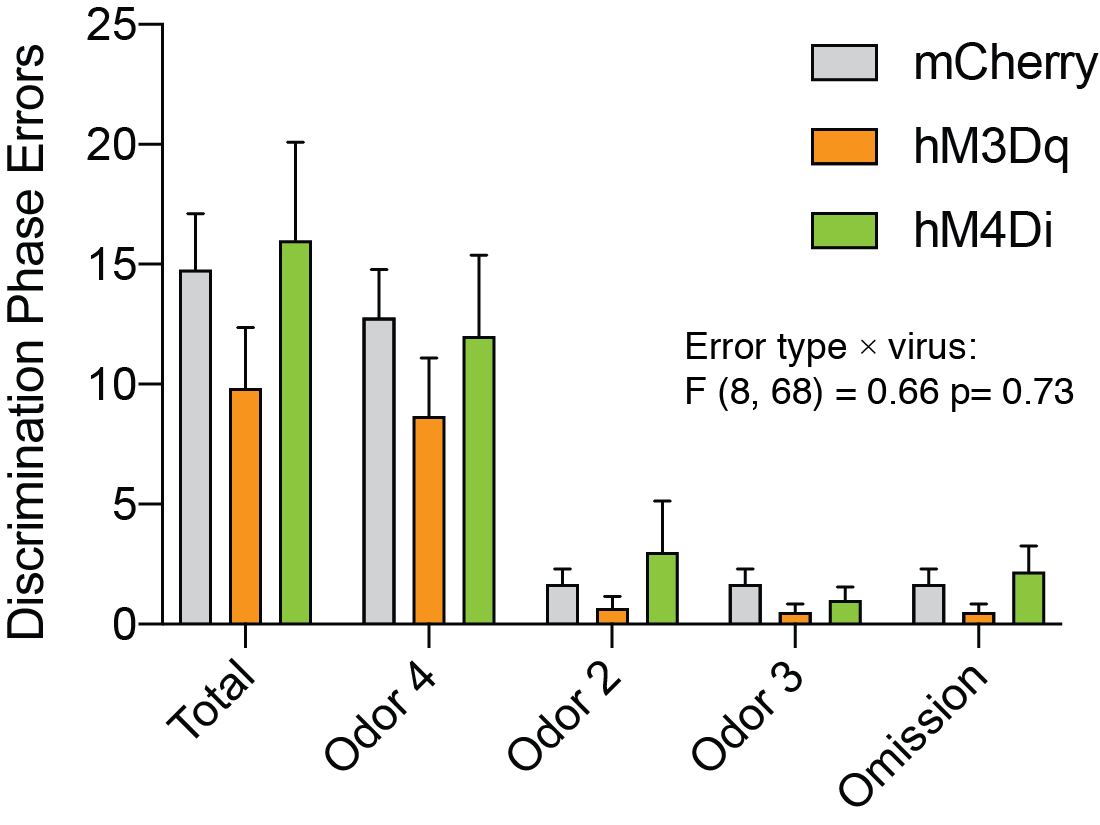

**Supplementary Figure 2.** Error type and virus do not significantly interact during discrimination phase, when all groups receive saline injections. p= 0.73, two-way repeated measures ANOVA; N= 9,6,5.

**Supplementary Table 1. RL model comparison for OVX and sham multiple choice reversal task data**

| Model | Learning rate α | Softmax β | Number of parameters per subject | Mean AIC |
| --- | --- | --- | --- | --- |
| ab | single | single | 2 | 152.40 |
| aab | per phase | single | 3 | 147.96 |
| abb | per phase | single | 3 | 149.31 |
| **aabb** | **per phase** | **per phase** | **4** | **144.92** |
